# C57 and DBA mouse strains express distinct cocaine avoidance phenotypes in an operant runway independent of differences in striatal dynorphin-enkephalin balance

**DOI:** 10.1101/2025.05.03.652074

**Authors:** Bailey C. Remmers, In Bae Choi, Kanako Matsumura, Amelia Nicot, Sydney Cox, Lauren K. Dobbs

## Abstract

**Rationale:** Human self-report data and rodent models indicate cocaine can induce negative affect, which may impact future cocaine use severity. Despite this, understanding of the neurobiology driving cocaine avoidance is limited. Within the striatum, the opioid peptides enkephalin and dynorphin are associated with cocaine reward and aversion, respectively. Additionally, striatal dynorphin signaling resulting from stress or cocaine withdrawal acts as a negative reinforcer to increase cocaine seeking.

**Objectives:** We validated the use of the cocaine self-administration runway in mice to measure the development of cocaine avoidance and tested whether mice with higher relative striatal dynorphin-to-enkephalin expression are resistant to developing cocaine avoidance.

**Methods:** Cocaine avoidance in the self-administration runway was measured in two inbred mouse strains, C57BL/6J and DBA/2J, known to have opposite striatal dynorphin/enkephalin milieus, and in mice lacking enkephalin selectively from striatal medium spiny neurons (D2-*Penk*KO).

**Results:** Both inbred strains developed cocaine avoidance, though they expressed it differently. Across training, DBA/2J mice increased their latency to self-administer cocaine, and C57BL/6J increased the number of retreats away from the cocaine-paired goal box. D2-*Penk*KOs developed similar cocaine avoidance compared to littermate controls.

**Conclusions:** Mice develop avoidance to self-administer cocaine in the runway across a range of strains. Pre-existing strain differences in the striatal dynorphin/enkephalin milieu, however, do not appear to alter the development of cocaine avoidance, and striatal enkephalin is not necessary for the development of cocaine avoidance. This suggests higher relative striatal dynorphin does not facilitate cocaine seeking by mitigating cocaine avoidance.

## Introduction

Cocaine is a psychostimulant drug with high addictive potential, and data from the National Survey on Drug Use and Health indicate an increased prevalence of cocaine use in recent years (Mustaquim et al. 2021; Orndorff et al. 2024). Despite this, there are no FDA-approved treatments for cocaine use disorder (Brandt et al. 2021). While the rewarding sensation and positive affective state elicited by cocaine contribute to its abuse potential, evidence from human self-report data indicates cocaine can also induce a negative affective state, which can affect the motivation to seek and take cocaine. Negative affect associated with acute cocaine administration is marked by feelings of anxiety, paranoia, and panic attacks (Anthony et al. 1989; Williamson et al. 1997; Kalayasiri et al. 2006). A similar negative affective state occurs during the “crash” as cocaine levels begin to decline, with the early phase of this crash characterized by agitation, nervousness, hypervigilance, and anorexia (Gawin and Kleber 1986; Kreek 1997). Together these data indicate that cocaine can induce opposing affective states, rewarding and aversive, and the culmination of these drive the motivational responses to either seek and take cocaine or to avoid it. Despite this knowledge, our understanding of the neural mechanisms underlying these opposing affective states and subsequent motivated behavior is limited.

Rodent models have been used for decades to probe the neurobiological mechanisms of cocaine reward. In addition, there is evidence that rodents experience cocaine’s negative affective state. Rats emit positive affect-related vocalizations during cocaine loading and when cocaine is at the subject’s preferred satiety level (Barker et al. 2014). In contrast, rats emit negative affect-related vocalizations when cocaine administration is clamped below the subject’s preferred satiety level or when forced to abstain (Barker et al. 2014; Klein et al. 2020). Place conditioning studies suggest these vocal calls may represent a negative motivational state because rats develop place avoidance to a context paired with a 15-minute withdrawal from a 0.75 mg/kg intravenous cocaine infusion (Ettenberg et al. 1999). Moreover, this negative affective state appears to drive the motivation to self-administer cocaine. Rats trained to traverse a straight alley runway to receive intravenous cocaine develop an approach-avoidance conflict over successive training trials (Ettenberg and Geist 1991; Geist and Ettenberg 1997; Ettenberg 2004; Jhou et al. 2013; Li et al. 2021; Parrilla-Carrero et al. 2021). Although rats volitionally culminate each session by reaching the goal box and receiving a cocaine infusion, they develop progressive cocaine avoidance characterized by an increasing number of retreats away from the goal box where cocaine is delivered (Ettenberg 2004). Additionally, pretreatment with the anxiolytic drug diazepam reduces retreats in this procedure (Ettenberg and Geist 1991), suggesting that avoidance in this task represents a conditioned anxiety-like state as a result of cocaine self-administration.

While these studies suggest cocaine avoidance arises from the negative, anxiogenic effects associated with the cocaine “crash”, acute cocaine administration can induce a distinct aversive state. Feelings of paranoia are reported to peak at the height cocaine administration during a two-hour cocaine binge, which then abated once access to cocaine ended (Kalayasiri et al. 2006). Consistent with this, plasma corticosterone levels are increased in rats following acute or repeated cocaine (Yang et al. 1992), and acute cocaine decreases the time spent in the open arms of an elevated plus maze (Yang et al. 1992; Rogerio and Takahashi 1992; Schank et al. 2008). This acute cocaine-induced anxiety state may contribute to cocaine avoidance. We recently used a trace conditioning procedure to show that the motivational state immediately following cocaine administration can generate place avoidance (Nicot et al. 2025). In this procedure, cocaine was administered immediately after removal from the conditioning apparatus, thereby imbuing the preceding context with cocaine’s transient aversive state. This procedure has been shown to condition place avoidance to a range of abused drugs, including ethanol, nicotine, and amphetamine (Fudala and Iwamoto 1987, 1990; Cunningham and Okorn 1997). Thus far, evidence suggests cocaine induces opposing affective states that interact to drive motivated behavior to seek and take cocaine. However, the neurobiological mechanisms by which cocaine induces negative affect, be it due to acute cocaine administration or falling cocaine levels, is fairly limited compared to our understanding of cocaine’s rewarding effects.

Within the striatum, the activity of mu opioid receptors (MOR) and kappa opioid receptors (KOR) have emerged as important mediators of the rewarding and aversive aspects of cocaine, respectively. Pharmacological blockade of MORs in the ventral striatum attenuates acquisition and expression of cocaine place preference (Soderman and Unterwald 2008; Dai et al. 2022). Similarly, selective deletion of MORs from dopamine D2 receptor expressing medium spiny neurons (D2-MSN) impairs the acquisition of cocaine place preference (Remmers et al. 2025). MOR-mediated facilitation of cocaine reward appears due to the opioid peptide enkephalin, as withdrawal from repeated cocaine increases expression of the enkephalin gene, *Penk*, in the striatum and increasing the tone of met-enkephalin in the ventral striatum potentiates the acquisition of cocaine preference (Dai et al. 2022).

In contrast to met-enkephalin/MOR-mediated effects, dynorphin-mediated KOR activation facilitates the negative affective state associated with cocaine withdrawal. Repeated cocaine administration increases striatal expression of the prodynorphin (*Pdyn*) gene (Steiner and Gerfen 1993), which is thought to generate an aversive stress-response because ventral striatal KOR signaling is acutely aversive in itself (Al-Hasani et al., 2015; Pirino et al., 2020). Moreover, KOR agonists and stress-induced dynorphin release reinstate operant cocaine seeking after extinction (Beardsley et al. 2005; Carey et al. 2007; Valdez et al. 2007; Redila and Chavkin 2008), suggesting dynorphin-mediated KOR signaling during cocaine withdrawal increases cocaine seeking via negative reinforcement. Conversely, evidence from trace conditioning indicates that low ventral striatal dynorphin levels are associated with the transient aversive state following cocaine administration. Mice that developed cocaine place avoidance in trace conditioning had lower ventral striatal *Pdyn* levels compared to mice that developed cocaine preference or remained neutral. Accordingly, mice with pre-existing higher striatal dynorphin relative to enkephalin were resistant to developing cocaine place avoidance, suggesting high striatal dynorphin may unmask the pro-motivational drive to seek cocaine (Nicot et al. 2025).

Taken together, these data suggest that the balance between striatal enkephalin and dynorphin may be an important mechanism for maintaining the motivational drive to seek and take cocaine. Because the approach-avoidance conflict to self-administer cocaine in the runway has not yet been published in mice, we first validated this procedure in a mouse model using two inbred mouse strains: DBA/2J and C57BL/6J. We next investigated how enkephalin-dynorphin balance within the striatum regulates the development of cocaine avoidance. We predicted that mice with a naturally higher dynorphin-to-enkephalin milieu in the striatum would show less cocaine avoidance. To address this, we compared cocaine avoidance behaviors between C57BL/6J and DBA/2J mice, which are known to have opposite striatal expression of enkephalin and dynorphin. DBA/2J mice have more striatal *Pdyn* expression, dynorphin A 1-13, dynorphin A 1-8, and KOR binding sites than C57BL/6J mice (Jamensky and Gianoulakis 1997). Additionally, DBA/2J have less *Penk* expression (Jamensky and Gianoulakis 1999) and binding sites for its principal receptor, the delta opioid receptor (de Waele and Gianoulakis 1997). Finally, we tested transgenic mice lacking enkephalin selectively from striatal D2-MSNs (D2-*Penk*KO) to directly interrogate how the striatal enkephalin/dynorphin balance contributes to cocaine avoidance.

## Methods

### Animals

Male and female mice between 8-26 weeks old were used for all experiments. All mice were housed in a temperature and humidity-controlled environment under a reversed 12:12 hour light cycle (lights on at 9:30 am), and experiments were run in the subject’s dark cycle. Animals had *ad libitum* access to food and water during recovery from surgery, and a subset of mice were mildly food-restricted at the beginning of training to encourage exploration of the runway (see *Statistics*). Animals were individually housed beginning one week before testing and throughout the experiment. All procedures were approved by the Institutional Animal Care and Use Committees of the University of Texas at Austin. DBA/2J (RRID: IMSR_JAX:000671) and C57BL/6J (RRID: IMSR_JAX:000664) mice were obtained from Jackson Labs. We generated mice with a selective loss of enkephalin from striatal D2-MSNs (D2-*Penk*KO) by crossing *Adora2a-Cre^+/-^* mice (B6.FVB(Cg)-Tg(Adora2a-cre)KG139Gsat/Mmucd, MMRC #36158) with mice possessing a floxed *Penk* gene (Schurmann, 2009). *Adora2a-Cre^+/-^* mice are congenic on a C57BL/6J background and the resulting D2-*Penk*KO and *Penk^f/f^* littermate controls mice have been crossed onto the *Adora2a-Cre^+/-^* line for more than 10 generations. These mice are bred in house and genotyped with Transnetyx. We previously showed that these D2-*Penk*KO mice lack *Penk*, as well as met-enkephalin and leu-enkephalin, selectively from striatal D2-MSNs (Matsumura et al. 2023).

### Surgery

Mice were deeply anesthetized with isoflurane and shaved over the mid-scapular region and the ventral right side of the neck. The right jugular vein was isolated and a catheter (Instech) was inserted through a small hole in the vein made using vannas spring scissors and secured in place with suture thread. The catheter was fed subcutaneously and attached to a single channel vascular access button (Instech) which was positioned in the mid-scapular region. Mice were treated post-operatively with daily antibiotic (cefazolin, 20 mg/kg, IV) and analgesic (carprofen, 10 mg/kg, IP) over a 5-day recovery period before testing began. Catheter patency was confirmed by immediate loss of the righting reflex following administration of sodium brevital (10 mg/kg; IV). Mice that failed patency were removed from the study (n = 6). Mice received heparinized saline (30 U/mL; 0.02 mL, IV) daily to maintain patency.

### Drugs

Cocaine hydrochloride (Sigma) was dissolved in saline and self-administered intravenously (1.0 or 1.5 mg/kg/infusion) during runway training or experimenter delivered as a pre-session challenge (1.5 mg/kg, IV). The volume of injection was based on animal weight. Morphine sulfate salt pentahydrate (Spectrum Chemical) was dissolved in saline and administered on challenge days 20 minutes before testing (3 mg/kg, IP).

### Runway Apparatus

Design of the approach-avoidance runway was based on the original rat runway designed by Dr. Ettenberg and colleagues and scaled down to use with mice (Geist and Ettenberg 1990). The straight alley runway (5 ft x 6 in) was constructed from grey acrylic and flanked by an equally sized “start box” and “goal box” on either end. The goal box was distinguished from the start box by a textured floor and black and white striped walls. The start box and goal box were equipped with removable guillotine doors in order to confine the mouse in either box. A magnetic track was positioned above the runway to float the drug swivel and infusion line over the runway, which allowed subjects to freely move in the apparatus. The track consisted of parallel bar magnets running down the length of the maze. A pot magnet of opposite polarity floated between the bars and housed a single-channel swivel. Because mice traverse the runway very quickly at baseline, we installed hurdles (4 cm high) in the maze spaced 10 cm apart to have a better dynamic range for baseline goal box latency. Mouse position was tracked by 14 infrared beams spaced 7.62 cm apart controlled by custom software (LabView, National Instruments). Cocaine was delivered via an infusion pump (Med Associates) controlled by custom-written software (LabView, National Instruments).

### Cocaine Runway Procedure

#### Initial Training

Male and female C57BL/6J, DBA/2J, D2-*Penk*KO, *Penk*^f/f^ littermate controls were trained to self-administer cocaine in the runway apparatus. Mice were first habituated to the apparatus and handling in one single 15-minute session, in which they were able to freely traverse the start box and the alleyway but not enter the goal box. After habituation, mice received 10 cocaine self-administration training trials, with 2 trials per day over 5 days. Morning trials began as soon as the dark cycle started, and afternoon trials began 3 hours afterwards. On each training day, subjects were placed into the start box with the guillotine door closed for 1 minute, after which time the door was removed and the mouse was given 10 minutes to reach the goal box. Upon entering the goal box, the goal box door was closed, cocaine was automatically infused (1 mg/kg or 1.5 mg/kg), and subjects remained confined for 5 minutes. If the mouse did not reach the goal box within the time limit, they were gently shuttled to the goal box, cocaine was administered, and they were confined for 5 minutes. If mice did not reach the goal box on their own by the second trial, they were removed from the study. The subject’s position was tracked by infrared beams. After completing 10 cocaine self-administration training trials, subjects were tested with different drug challenges to assess the ability of other drugs to alter the baseline approach-avoidance behavior (described below).

#### Drug Challenges

To determine how acute administration of morphine or cocaine affects the developed avoidance to self-administer cocaine, we administered a morphine pretreatment (3 mg/kg, IP) or a cocaine pretreatment (1.5 mg/kg, IV) in separate, within subjects challenge tests. After completing 10 training trials in the cocaine runway as described above, DBA/2J mice, D2-*Penk*KO and *Penk*^f/f^ controls were challenged with morphine. In addition, D2-*Penk*KOs and *Penk*^f/f^ controls receiving an intervening cocaine challenge between initial training and the morphine challenge.

Morphine challenge was administered in the subject’s home cage 20 minutes before being placed into the start box. Subjects were then allowed to traverse the runway as in a normal training trial where upon reaching the goal box 1.5 mg/kg cocaine (IV) was administered and subjects were confined for 5 minutes. Thus, subjects had the subjective experience of morphine in combination with cocaine while in the goal box. To assess how the reinforcing value of the goal box had been changed after this morphine challenge, on the following day, subjects received an IP injection of saline in their home cage 20 minutes before being placed in the start box. They were then allowed to traverse the runway and received cocaine (1.5 mg/kg, IV) upon entering the goal box.

Cocaine challenge (1.5 mg/kg) was intravenously delivered in the start box. One minute after the cocaine infusion, subjects were allowed to traverse the runway. Upon entering the goal box, subjects received an additional cocaine infusion (1.5 mg/kg, IV) and were confined for 5 minutes.

### Food Runway Procedure

Subjects were habituated to 2.0 g crushed Reese’s Pieces treats (approximately 3 treats) in their home cage over 3 days before the start of the experiment. Subjects underwent a habituation trial followed by training trials, which had the same schedule and timing (2 trials/day, AM trials beginning at the start of the dark phase, separated by 3 hours) as in the cocaine runway. In the food runway, though, subjects received access to 1 crushed Reese’s Piece (0.6 – 0.8 g) in a weigh boat upon entering the goal box and were confined to the goal box for 3 minutes to consume the treat. Across the training trials, treat consumption was estimated by weighing the food before and after each trial. We observed that subjects reliably consumed the candy filling (0.05 ± 0.007 g) while leaving the candy shell uneaten.

### Statistics

Mouse position in the runway was tracked using infrared beams with custom-written software (LabVIEW, National Instruments), and each beam break was timestamped to determine the subject’s position over time. From these timestamps, we calculated the following dependent variables: total distance traveled, latency to exit the start box, latency to reach the goal box, number of retreat bouts, distance spent retreating, and average distance per retreat bout. A retreat bout was defined as continuous movement away from the start box so that the mouse had to break at least 2 consecutive infrared beams in the direction toward the start box.

DBA/2J and C57BL/6J training data sets were analyzed with sex as a biological variable using a repeated measures three-way ANOVA (sex x reinforcer x trial). If the main effect of sex or interactions with sex were not significant, data were collapsed across sex and analyzed using repeated measures two-way ANOVA (reinforcer x trial). Sample sizes per sex in the D2-*Penk*KO experiment and cocaine challenges were too small (e.g., ≤ 2) to permit analysis with sex as a statistical factor, though individual data are presented labeled by sex. Thus, the D2-*Penk*KO/*Penk*^f/f^ training data sets were analyzed using a three-way repeated measures ANVOA (genotype x reinforcer x trial). Additionally, during training a subset of subjects was mildly food-restricted to encourage exploration. However, we found no effect of food restriction on the latency measures or retreat variables, thus this was not included as a factor in the analyses. Data that were averaged across the last five training trials were compared between reinforcer groups or strains using independent samples t-tests or Mann-Whitney U tests if the data sets were not normally distributed. Cocaine challenge data for D2-*Penk*KO and *Penk*^f/f^ mice were compared to baseline using a two-way repeated measures ANOVA (genotype x day). Morphine challenge data were analyzed using one-way repeated measures ANOVA for DBA/2J mice and a two-way repeated measured ANOVA for D2-*Penk*KO and *Penk*^f/f^ mice (genotype x day). Multifactorial data sets with missing values were analyzed using a Mixed Effects Model and violations to sphericity in repeated measures analyses were corrected using the Greenhouse-Geiser method. Significant interactions from ANOVAs were followed up with Šidák-corrected t-test comparisons. All data were analyzed using Prism (GraphPad) and results were considered significant at an alpha of 0.05. Data are presented as mean ± SEM or individual data points are labeled by sex.

## Results

### DBA/2J mice develop avoidance to self-administer cocaine in the runway

We tested cocaine approach-avoidance in DBA/2J mice using a scaled-down version of the rat runway created by Ettenberg and colleagues (Ettenberg and Geist 1991) (Fig. 1a). Approach-avoidance was determined by the change in several within-subjects dependent variables across training trials, including the latency to exit the start box, latency to the goal box, number of retreats away from the goal box, total retreat distance, and average distance per retreat. Consistent with prior literature in rats, cocaine-trained DBA/2J mice developed avoidance behaviors across training as well as compared to food-trained mice (Fig. 1b). Cocaine-trained mice developed longer latencies to exit the start box over training compared to food-trained mice (Trial x Reinforcer, F_9,239_ = 2.37, *p* < 0.05), and this was different specifically at trials 8 (t_266_ = 4.16, *p* < 0.001) and 10 (t_266_ = 2.89, *p* < 0.05; Fig. 1c-e). Both reinforcer groups decreased the time it took to reach the goal box across training trials (Trial, F_3.2,78.48_= 10.45, *p* < 0.0001); however, cocaine-trained mice stabilized at longer latencies compared to food-trained mice (Trial x Reinforcer, F_9,221_ = 2.82, *p* < 0.01; Fig. 1f-h). Goal box latency for cocaine-trained mice plateaued by trial 3, while food-trained mice continued to decrease their time to goal (post hoc between reinforcers for trials 3 – 10, respectively: t_15_ = 3.28, *p* < 0.05; t_15_ = 3.71, *p* < 0.05; t_14_ = 3.44, *p* < 0.05; t_15_ = 3.62, *p* < 0.05; t_15_ = 3.51, *p* < 0.05; t_15_ = 3.58, *p* < 0.05; t_15_ = 3.63, *p* < 0.05; t_15_ = 3.54, *p* < 0.05). This was supported by additional comparison of the average of the last 5 trials (Mann-Whitney U: 35, *p* < 0.01). Additionally, males took longer to reach the goal box compared to females, regardless of reinforcer (Trial x Sex, F_9,221_ = 3.27, *p* < 0.001); however, post hoc comparisons indicated nominally greater latency only at trial 2 (t_23.68_ = 2.99, *p* = 0.06).

**Fig. 1.**
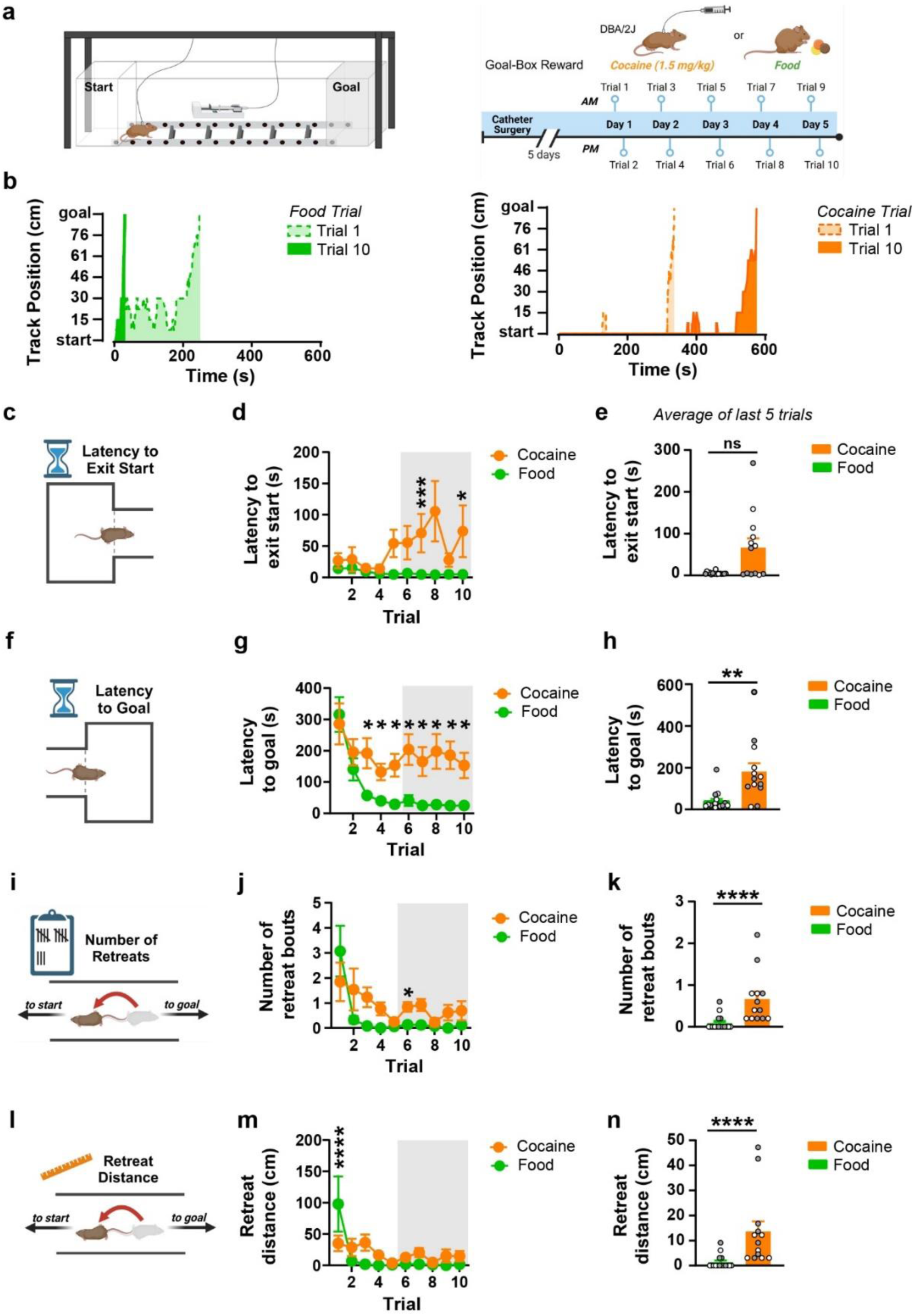
DBA/2J mice develop avoidance to self-administer cocaine in the runway. **a)** Schematic of the approach-avoidance runway (left) and experimental timeline of cocaine or food training (right). **b)** The track position of representative food-trained (left, green) and cocaine-trained (right, orange, 1.5 mg/kg) mice during training trial 1 (dashed line) and trial 10 (solid line). **c-e)** Cocaine-trained mice (orange, n = 13) developed greater latency to exit the start box, specifically at trials 7 and 10 compared to food-trained mice (green, n = 16) (d). The average latency to exit the start box over the last 5 trials was greater for cocaine-trained than food-trained mice (e). **f-h)** Cocaine-trained mice had higher latency to reach the goal box than food-trained mice between trials 3-10 (g), and when averaged over the last 5 days of training (h). **i-j)** The number of retreats decreased for both reinforcer groups, but cocaine-trained mice had more retreats on trial 6 (j). Averaged retreats over the last 5 training days were higher for cocaine-trained than food-trained mice (k). **l-n)** Retreat distance was higher for food-trained mice on trial 1 (m), but cocaine-trained mice had higher average retreat distances on the last 5 training days (n). ns = not significant, * *p* < 0.05, ** *p* < 0.01, *** *p* < 0.001, **** *p* < 0.0001. Data are shown as mean ± SEM. Grey shaded box in panels d, g, j, and m indicates the averaged trials shown in the corresponding bar graphs. Individual values are shown as male (grey) and female (white). Schematics created with Biorender.com.

Retreat variables were also different between food-trained and cocaine-trained DBA/2J mice. Across training trials, and regardless of reinforcer, subjects progressively decreased the number of retreats (Trial, F_2.1,48.62_ = 8.16, *p* < 0.001; Fig. 1i-k), total retreat distance (Trial, F_1.63,37.58_ = 7.2, *p* < 0.01; Fig. 1l-n), and distance per retreat (Trial, F_1.08,25.03_ = 6.33, *p* < 0.05; Supplemental Fig. 1c). Similar to the latency to goal, though, cocaine-trained mice plateaued at slightly more retreats later in training (Trial x Reinforcer, F_9,208_ = 2.38, *p* < 0.05), specifically at trial 6 (t_17.28_ = 3.37, *p* < 0.05), and with a trend at trial 7 (t_15.06_ = 3.14, *p* = 0.06). There were also differences between reinforcer groups in the total retreat distance (Trial x Reinforcer, F_9, 208_ = 3.75, *p* < 0.001) and distance traveled per retreat bout (Trial x Reinforcer, F_9, 208_ = 5.15, *p* < 0.0001); however, post hoc analysis did not reveal specific trials where cocaine-trained mice were greater than food-trained mice. Instead, food-trained mice showed greater retreat distance (post hoc, t_253_ = 3.75, *p* < 0.01) and distance traveled per retreat bout (post hoc, t_253_ = 5.38, *p* < 0.0001) on the first trial. This was due to food-trained males having larger retreat-related variables on the first trial relative to cocaine-trained males or food-trained females (see Table 1 for sex effect statistics). Since sex differences were not apparent for any retreat variable after the first trial, we collapsed across sex to compare the average of the last 5 trials between reinforcers. This confirmed cocaine-trained mice had more retreats (Mann-Whitney U: 18.5, *p* < 0.0001) and greater retreat distance (Mann-Whitney U: 16, *p* < 0.0001). Notably, though, retreats were fairly low (cocaine: 0.71 ± 0.17; food: 0.09 ± 0.04), as was the distance spent retreating (cocaine: 15.02 ± 4.0 cm; food: 1.33 ± 0.68 cm), potentially indicating that retreat behavior is not the most robust measure of cocaine avoidance for the DBA/2J mouse strain. General locomotor activity was different between reinforcer groups (Trial x Reinforcer: F_9,206_ = 4.87, *p* < 0.0001) and sexes (Trial x Sex: F_9,206_ = 2.47, *p* < 0.05) across trials (Supplemental Figure 1a). However, post hoc follow up did not identify differences between reinforcer groups at any trial. Additionally, post hoc comparisons indicated males had higher locomotion than females at trial 2 (t_16.14_ = 3.79, *p* < 0.05). Thus, the development of cocaine avoidance in DBA/2J mice does not appear to be due to differences in general locomotion.

**Table 1.**
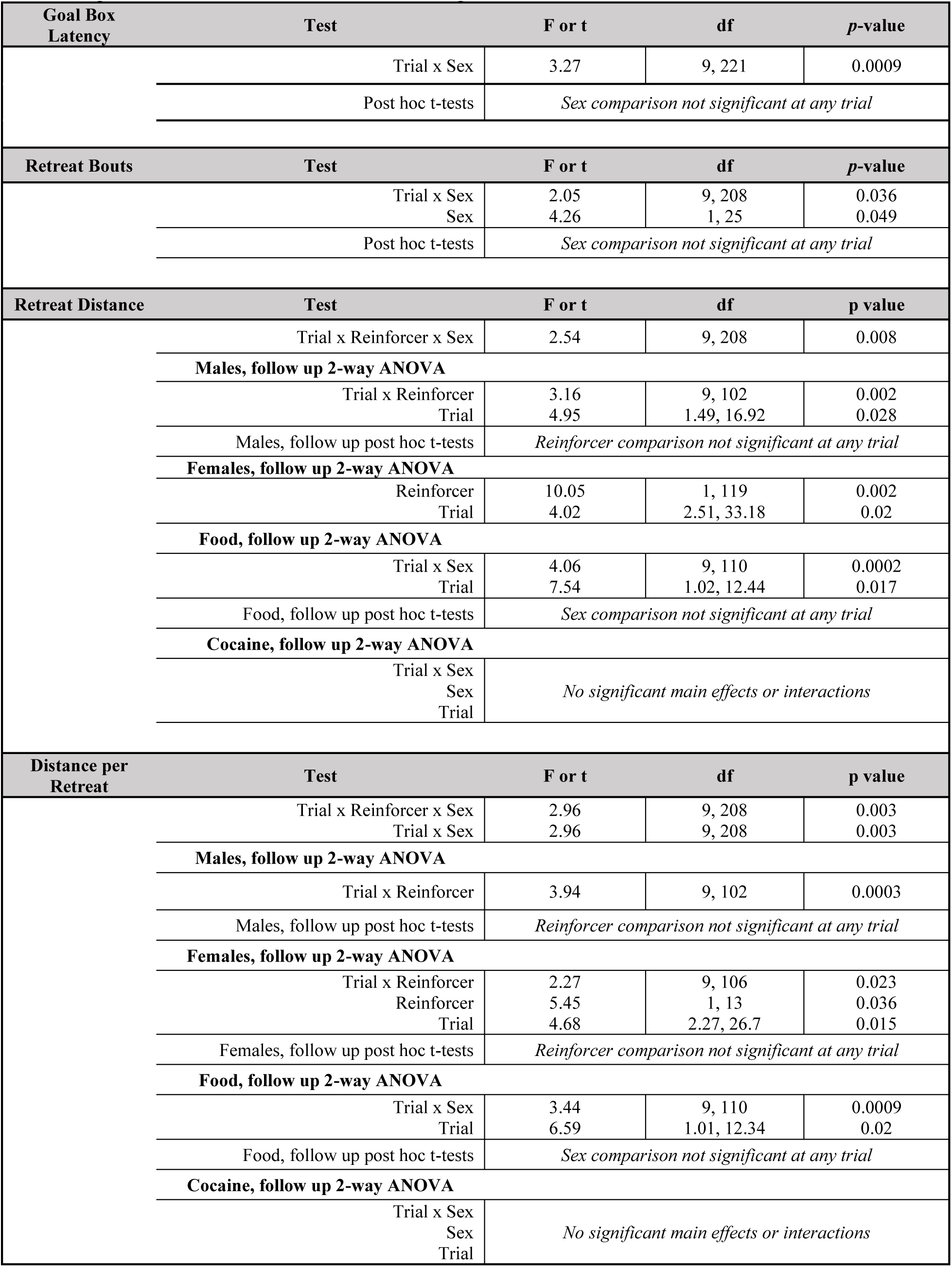
Significant sex effects results for DBA/2J training trials.

### C57BL/6J mice develop avoidance to self-administer cocaine in the runway

We next sought to determine whether a different inbred mouse strain, the C57BL/6J, develops cocaine avoidance in the self-administration runway (Fig. 2a). Similar to the DBA/2J findings, cocaine-trained C57BL/6J mice developed signs of avoidance (more retreats and longer latencies) over training compared to their food-trained counterparts (Fig. 2b), though no effects of sex were observed across any of the variables. While there were no differences between reinforcer groups or across trials for the latency to exit the start box (Fig. 2c-e), cocaine-trained subjects took longer to reach the goal box compared to food-trained subjects (Reinforcer, F_1, 22_ = 5.30, *p* < 0.05; Fig. 2f-g). This was supported by comparison of the averaged latency across the last 5 trials (Mann-Whitney U: 11, *p* < 0.001, Fig. 2h). Cocaine-trained C57BL/6J mice increased their retreats across training trials, while food-trained mice maintained relatively no retreats across training (Trial x Reinforcer, F_9, 191_ = 2.35, *p* < 0.05; Fig. 2i-j). Post hoc analysis indicated a nominal difference between reinforcer groups at trial 5 (t_13_ = 3.02, *p* = 0.09) and a significant difference at trial 8 (t_13_ = 4.28, *p* < 0.01). Cocaine-trained C57BL/6J mice also had overall greater total retreat distance (Reinforcer, F_1, 22_ = 11.13, *p* < 0.01; Fig. 2l-m) and distance per retreat (Reinforcer, F_1, 22_ = 11.19, *p* < 0.01; Supplemental Fig. 1d) than food-trained counterparts. Analysis of the averaged last 5 trials supported these differences for the retreats (Mann-Whitney U: 16, *p* < 0.001, Fig. 2k) and retreat distance (Mann-Whitney U: 13, *p* < 0.001, Fig. 2n). The increased retreats and retreat distance were also consistent with greater overall distance travelled in cocaine-trained versus food-trained mice (Reinforcer, F_1,22_ = 9.8, *p* < 0.01; Supplemental Fig. 1b). Taken together, these results indicate that C57BL/6J mice develop avoidance to self-administer cocaine in a runway procedure.

**Fig. 2.**
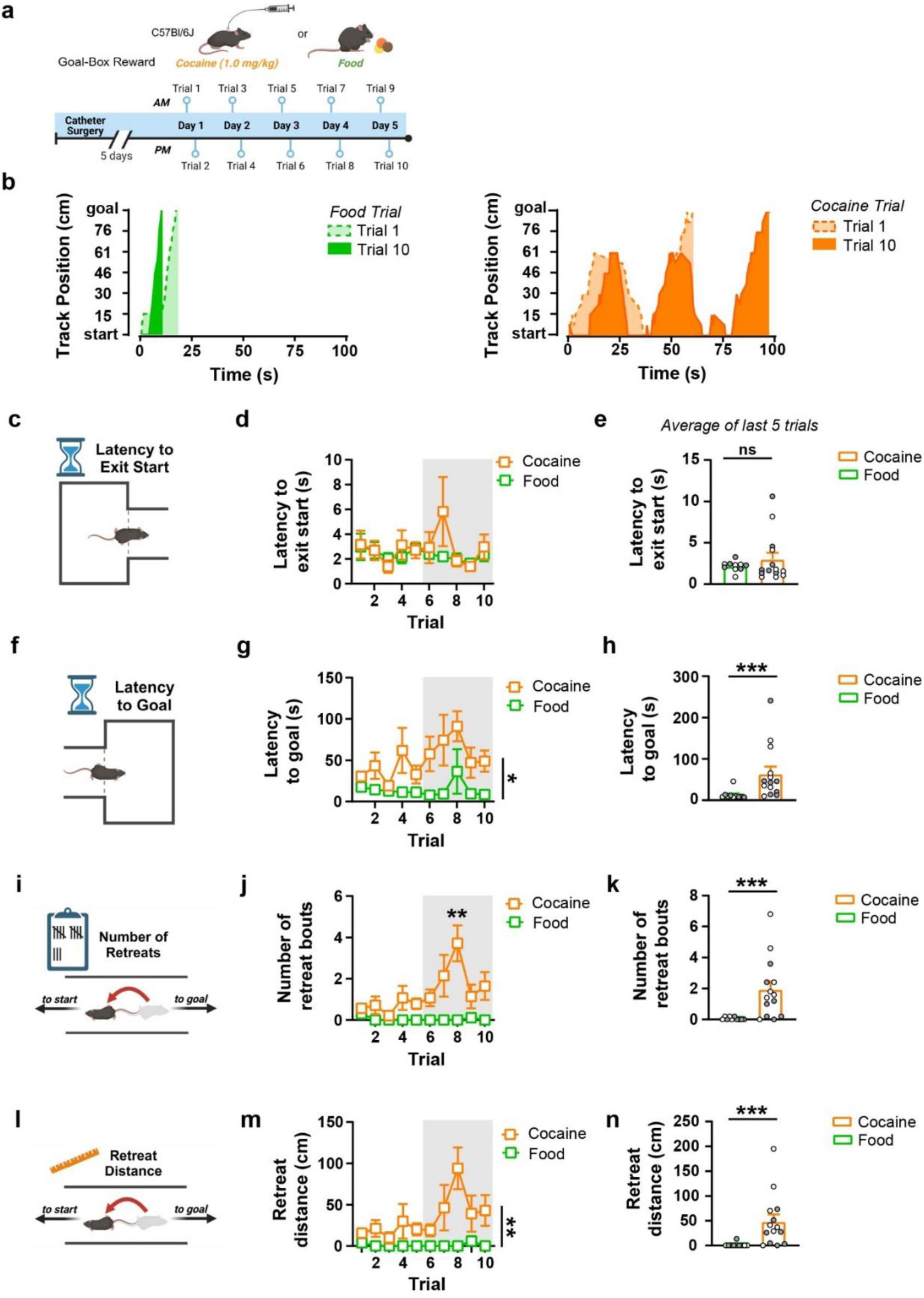
C57BL/6J mice develop avoidance to self-administer cocaine in the runway. **a)** Experimental timeline of cocaine or food training. **b)** The track position of representative food-trained (left, green) and cocaine-trained (right, orange) mice during training trial 1 (dashed line) and trial 10 (solid line) is shown. **c-e)** Latency to exit the start box was similar between cocaine-trained (orange; n = 14) and food-trained (green; n = 10) mice across trials (d) and for the averaged last 5 training trials (e). **f-h)** Cocaine-trained mice took longer to reach the goal box compared to food-trained mice across all trials (g) and for averaged last 5 training trials (h). **i-j)** The number of retreats increased over training trials for cocaine-trained mice and was higher than food-trained mice on trial 8 (j) and when comparing the averaged last 5 trials (k). **l-n)** Retreat distance was higher for cocaine-trained compared to food-trained mice across all trials (m) and for the averaged last 5 trials (n). ns = not significant, * *p* < 0.05, ** *p* < 0.01, *** *p* < 0.001. Data are shown as mean ± SEM. Grey shaded box in panels d, g, j, and m indicates the averaged trials shown in the corresponding bar graphs. Individual values are shown as male (grey) and female (white). Schematics created with Biorender.com.

### DBA/2J and C57BL/6J mouse strains express cocaine avoidance differently in the runway

Prior literature indicates high striatal dynorphin is associated with greater cocaine seeking (Beardsley et al. 2005; Carey et al. 2007; Valdez et al. 2007; Redila and Chavkin 2008) and resistance to developing cocaine place avoidance (Nicot et al. 2025). Based on this, we hypothesized that DBA/2J mice, which have higher striatal dynorphin relative to enkephalin, would be more resistant to developing cocaine avoidance in the self-administration runway. Although both inbred mouse strains developed signs of cocaine avoidance, the manner in which avoidance was expressed was different. DBA/2J mice had longer latencies to exit the start box (Strain, F_1, 25_ = 9.51, *p* < 0.01, Supplemental Fig. 2a) and to reach the goal box (Strain, F_1, 25_ = 19.07, *p* < 0.001, Supplemental Fig. 2b) compared to C57BL/6J. In contrast, C57BL/6J mice primarily expressed cocaine avoidance through the development of retreat behaviors. C57BL/6J mice initially expressed very few retreats and increased retreats over training, while DBAs reduced their retreat behavior over training (retreat bouts: Strain x Trial, F_9, 233_ = 4.77, *p* < 0.0001; retreat distance: Strain x Trial, F_9, 224_ = 3.1, *p* < 0.01; Supplemental Fig. 2c-d). Post hoc analyses indicated C57BL/6J showed more retreats (t_13.92_ = 3.89, *p* < 0.05) and greater retreat distance (t_13.53_ = 3.51, *p* < 0.05) specifically at trial 8. Similarly, C57BL/6J mice showed greater overall distance per retreat compared to DBA/2J (Strain, F_1,279_ = 8.79, *p* < 0.01; Supplemental Fig. 2e), which likely contributed to the increase in locomotion at trial 8 (Strain x Trial, F_9, 221_ = 3.42, *p* < 0.001; t_14.47_ = 3.37, *p* < 0.05; Supplemental Fig. 2f).

### Deletion of striatal enkephalin does not alter the development of cocaine avoidance

Although DBA/2J and C57BL/6J mice have opposite enkephalin/dynorphin milieus, these strains also differ in other ways. Therefore, to directly investigate how a shifted balance between striatal enkephalin and dynorphin affects the development of cocaine avoidance in the self-administration runway, we used a transgenic mouse line with a selective deletion of *Penk* from striatal D2-MSNs (D2-*Penk*KO). Although some measures indicated D2-*Penk*KOs may have slightly less pronounced cocaine avoidance compared to *Penk^f^*^/f^ littermate controls, overall, both genotypes developed avoidance to self-administer cocaine, but not food (Fig. 3a). The latency to exit the start box was higher for cocaine-trained mice compared to food-trained (Reinforcer, F_1,31_ = 5.8, *p* < 0.05). While this effect seemed to develop over time and was more apparent in *Penk*^f/f^ controls, the interaction with genotype and trial only approached significance (Genotype x Reinforcer x Trial, F_9, 261_ = 1.83, *p* = 0.06; Fig. 3b-d). Similarly, latency to reach the goal was higher for cocaine-trained than food-trained mice (Reinforcer x Trial, F_9, 332_ = 2.83, *p* < 0.01; Fig. 3e-g). Post hoc comparisons indicated differences at trials 2 through 10 (t_391_ = 4.58, *p* < 0.0001; t_391_ = 4.78, *p* < 0.0001; t_391_ = 5.62, *p* < 0.0001; t_391_ = 5.26, *p* < 0.0001; t_391_ = 4.72, *p* < 0.0001; t_391_ = 3.63, *p* < 0.01; t_391_ = 3.42, *p* < 0.01; t_391_ = 5.57, *p* < 0.0001; t_391_ = 6.05, *p* < 0.0001). In addition, while D2-*Penk*KO mice had a slightly shorter goal box latency and the mean latency increased over time for *Penk*^f/f^ mice, this interaction only approached significance (Trial x Genotype, F_9, 332_ = 1.74, *p* = 0.07). Both genotypes also developed similar retreat behaviors for cocaine across training (Reinforcer x Trial, F_9, 331_ = 4.87, *p* < 0.0001) relative to their food-trained counterparts (Fig. 3h-j). Post hoc analysis indicated mice developed significantly more retreats by trial 8 (t_390_ = 3.8, *p* < 0.01) and this persisted through trials 9 and 10 (t_390_ = 5.74, *p* < 0.0001; t_390_ = 6.69, *p* < 0.0001). Likewise, the total retreat distance for cocaine-trained mice increased across trials for both genotypes compared to food-trained mice (Reinforcer x Trial, F_9, 330_ = 3.00, *p* < 0.01; Fig. 3k-m). Follow up analysis showed this was significant at trials 9 (t_389_ = 4.16, *p* < 0.001) and 10 (t_389_ = 4.35, *p* < 0.001). The distance per retreat bout increased over trials (Trial, F_3.54, 99.4_ = 3.62, *p* < 0.05) but was not different between genotypes or reinforcers (Supplemental Fig. 1g-h). These changes in retreats were accompanied by increased general locomotion for cocaine-trained mice relative to food-trained counterparts (Reinforcer x Trial, F_1, 261_ = 2.00, *p* < 0.05), which became apparent at trials 9 (t_312_ = 4.23, *p* < 0.001) and 10 (t_312_ = 4.4, *p* < 0.001; Supplemental Fig. 1e-f).

**Fig. 3.**
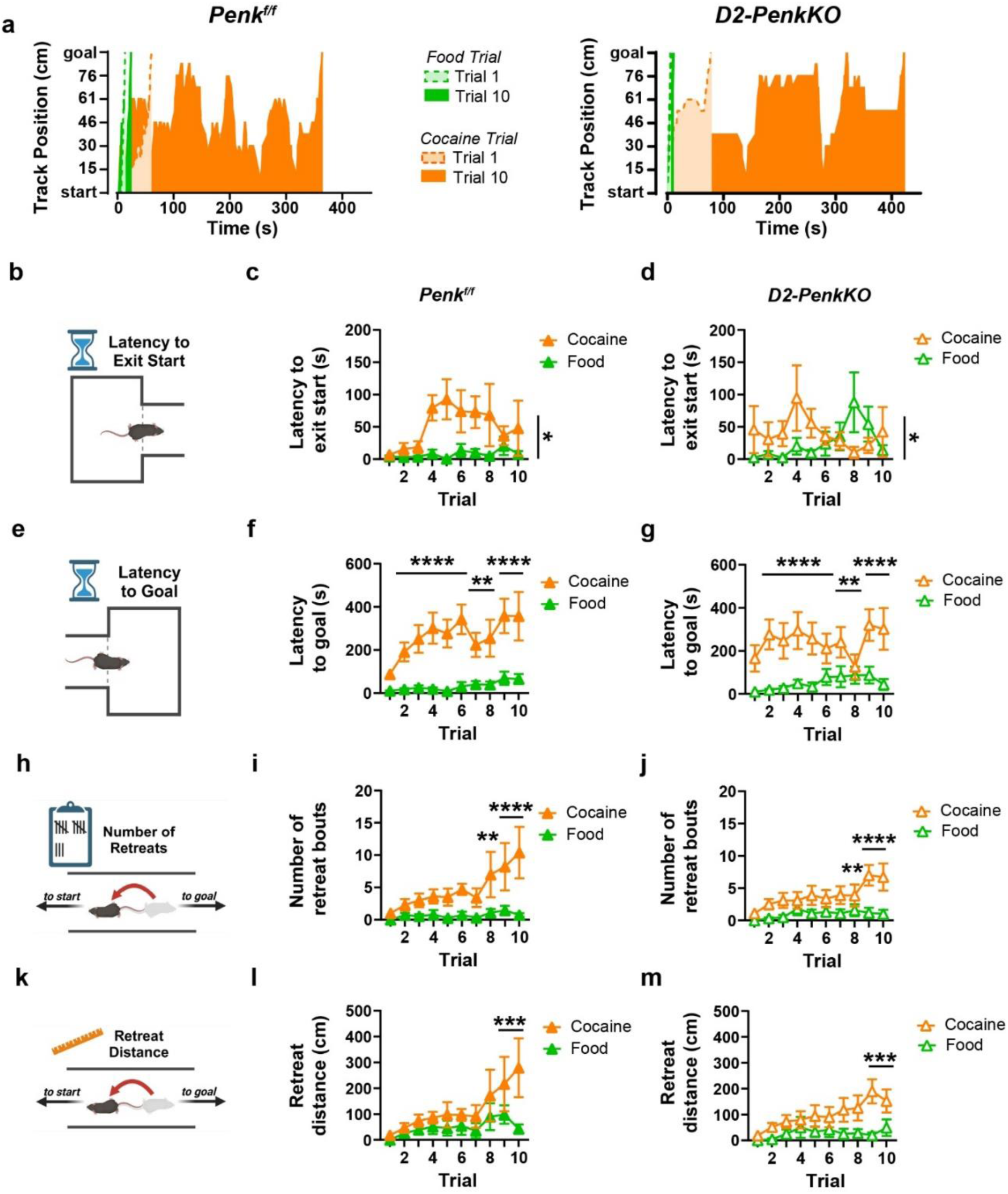
Lack of striatal enkephalin does not alter the development of cocaine avoidance. **a)** The track position of representative *Penk*^f/f^ controls (left) and mice lacking enkephalin from D2-MSNs (D2-*Penk*KO, right) is shown. Mice were trained to self-administer food (green) or cocaine (orange). Data are shown from training trial 1 (dashed line) and trial 10 (solid line). **b-d)** Latency to exit the start box was higher for cocaine-trained (orange; *Penk*^f/f^, n = 10; D2*Penk*KO, n = 10) compared to food-trained (green; *Penk*^f/f^, n = 9; D2*Penk*KO, n = 6) mice. This was true for both *Penk*^f/f^ controls (closed symbols, c) and D2-*Penk*KO mice (open symbols, d) as the Genotype x Reinforcer x Trial interaction only approached significance (*p* = 0.06). **e-g.** Cocaine-trained mice took longer to reach the goal box than food-trained mice from trials 2 – 10 for *Penk*^f/f^ controls (f) and D2-*Penk*KO mice (g). **h-j)** Cocaine-trained mice increased the number of retreats on trials 8 – 10 relative to food-trained mice for *Penk*^f/f^ controls (i) and D2-*Penk*KO mice (j). **k-m)** Cocaine-trained mice increased their retreat distance on trials 9 – 10 relative to food-trained mice for *Penk*^f/f^ controls (l) and D2-*Penk*KO mice (m). * *p* < 0.05, ** *p* < 0.01, *** *p* < 0.001, **** *p* < 0.0001. Data are shown as mean ± SEM. Schematics created with Biorender.com.

### A cocaine prime does not reduce avoidance to self-administer cocaine

Prior literature indicates that cocaine can induce a negative affective state following administration or withdrawal, and that this can influence the motivation to seek and take cocaine (Valdez et al. 2007; Nicot et al. 2025). However, the experience of a negative affective state associated with cocaine withdrawal while in the goal box may drive avoidance behaviors in the runway. Here, we tested the hypothesis that cocaine withdrawal contributed to the expression of cocaine avoidance in the runway. After completing 10 training trials in the runway, D2-*Penk*KO and *Penk*^f/f^ mice were challenged with an intravenous cocaine prime (1.5 mg/kg) while in the start box, and the subsequent approach-avoidance to the cocaine-associated goal box was measured. Compared to baseline values (average of trials 9 and 10), cocaine priming did not significantly reduce most of the dependent variables measured, including the latency to goal (Fig. 4d-f), number of retreats (Fig. 4g-i), retreat distance (Fig. 4j-l), distance per retreat (Fig. 4m-o), and the total distance traveled (Fig. S3a-b). Cocaine priming did, however, decrease the latency to exit the start box selectively in *Penk*^f/f^ controls (Genotype x Day, F_1, 10_ = 6.89, *p* < 0.05; post hoc, t_10_ = 3.46, *p* < 0.05; Fig. 4a-c). Together, these data suggest cocaine withdrawal may not be a large contributor to cocaine avoidance in the runway.

**Fig. 4.**
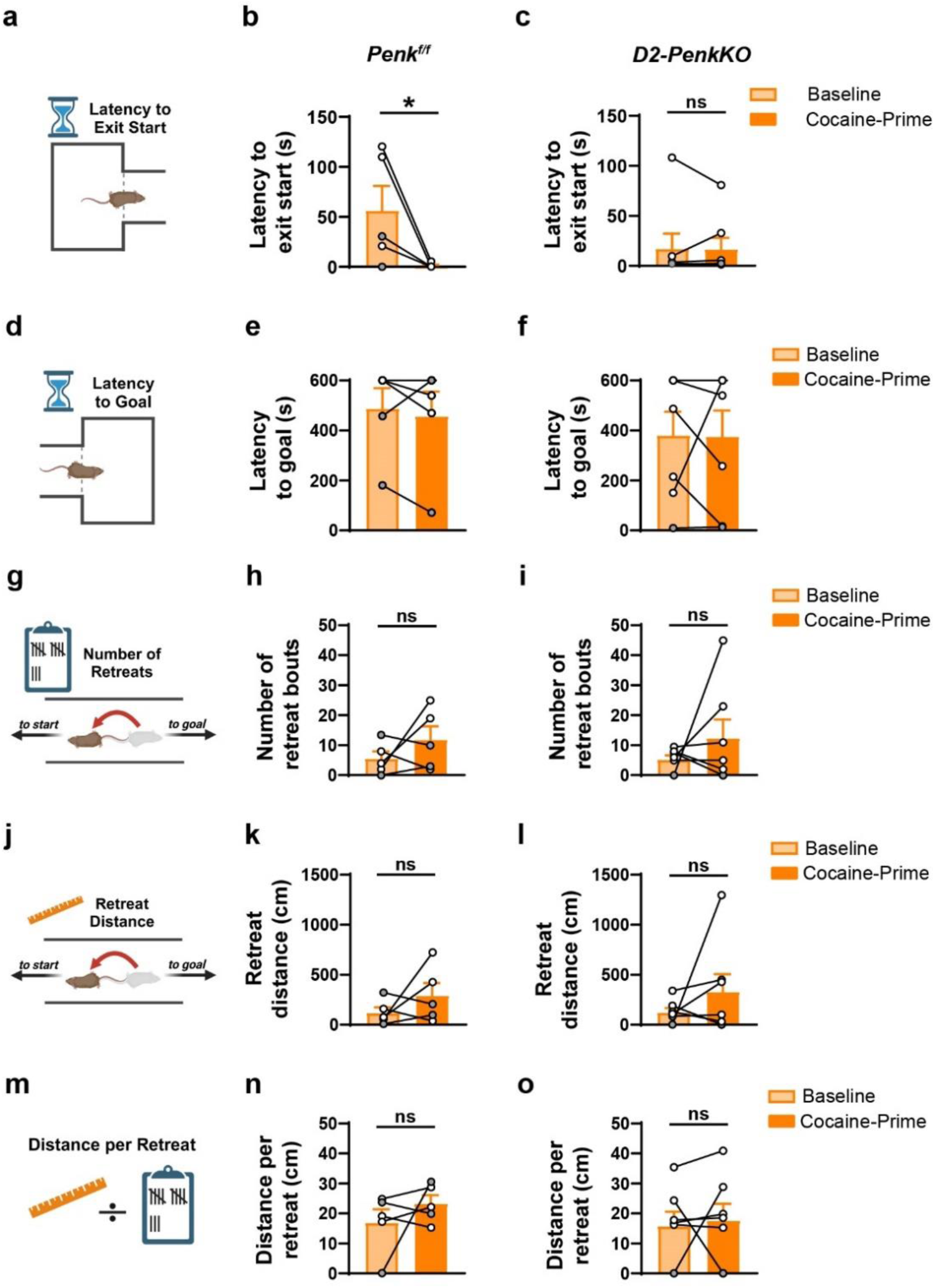
A cocaine prime does not reduce avoidance to self-administer cocaine. **a-c)** The latency to exit the start box decreased following administration of a cocaine prime in the start box (1.5 mg/kg, intravenous) compared to baseline (average over the last 2 training trials) for *Penk*^f/f^ mice (n = 5) (b) but not for mice lacking enkephalin from D2-MSNs (D2-*Penk*KO, n = 7) (c). **d-f)** The latency to reach the goal box was not affected by a cocaine prime for *Penk*^f/f^ (e) or D2-*Penk*KO (f) mice. **g-i)** Cocaine priming did not affect the number of retreats compared to baseline for *Penk*^f/f^ (h) or D2-*Penk*KO (i) mice. **j-l).** Retreat distance was not affected by a cocaine prime for *Penk*^f/f^ (k) or D2-*Penk*KO (l) mice. **m-o)** The average distance per retreat did not change following a cocaine prime in *Penk*^f/f^ (n) or D2-*Penk*KO (o) mice. ns = not significant, * *p* < 0.05. Data are shown as mean ± SEM. Individual values shown as male (grey) and female (white). Schematics created with Biorender.com.

### A morphine prime does not acutely reduce cocaine avoidance in the runway

Cocaine avoidance in the self-administration runway is reduced by pretreatment with the anxiolytic drug diazepam (Ettenberg and Geist 1991). In addition, rats show fewer retreats in the runway to self-administer a heroin-cocaine combination compared to cocaine alone (Guzman and Ettenberg 2004). Based on this, we hypothesized that a morphine prime would act similarly to benzodiazepines by alleviating cocaine-induced anxiety and decreasing cocaine avoidance in the runway. Twenty minutes after a morphine prime (3 mg/kg, IP), mice were allowed to traverse the runway and self-administer cocaine in the goal box just as in training trials. Morphine priming did not decrease the latency to exit the start box (Fig. 5a-d), retreat bouts (Fig. 5i-l), or retreat distance (Fig. 5m-p) in any strain tested, nor did we detect any sex differences in these variables. Surprisingly, though, morphine increased the time that D2-*Penk*KO and *Penk*^f/f^ mice took to reach the goal box (Day, F_1.82, 19.98_ = 6.15, *p* < 0.01; t_12_ = 3.3, *p* < 0.05; Fig. 5g-h). The latency to reach the goal also changed over the three days for DBA/2J mice (F_1.11, 13.3_ = 6.77, *p* < 0.05; Fig. 5f). Despite a larger proportion of DBA/2J mice increasing taking longer to reach the goal following a morphine prime (68.8%) than taking less time or showing no change (31.3%), follow up post hoc comparison between the morphine-primed day and baseline was not significant. General locomotion did not change after a morphine prime for DBA/2J, D2-*Penk*KO, or *Penk*^f/f^ mice, suggesting sedation or hyperactivity was not a confounding factor (Supplemental Fig. 3c-d); however, male DBA/2J mice were more active compared to females (F_1, 12_ = 6.71, *p* < 0.05).

**Fig. 5.**
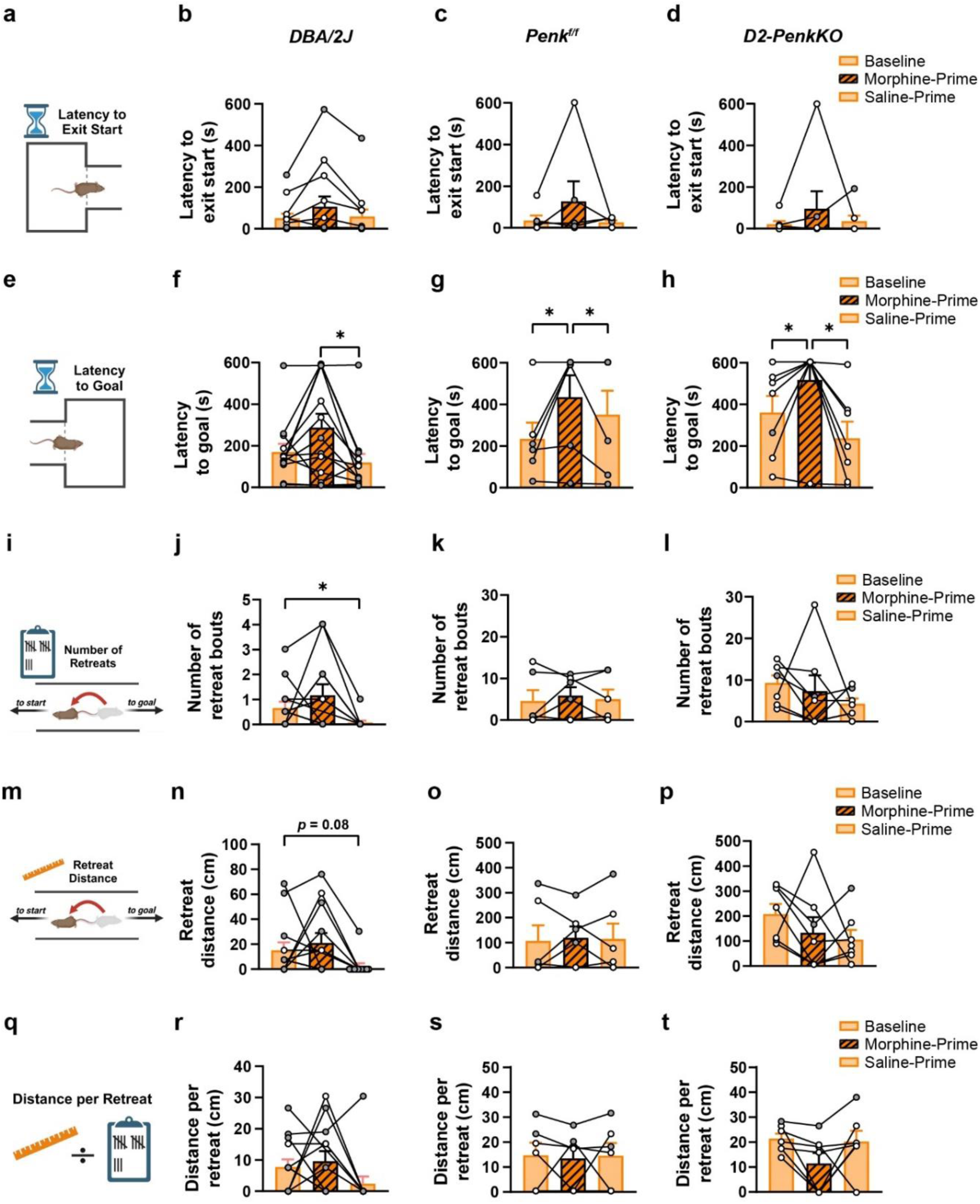
A morphine prime does not acutely reduce cocaine avoidance in the runway. **a-d)** Compared to baseline (average over last 2 training trials; left orange bar,), the latency to exit the start box was not affected by a morphine prime (3 mg/kg, IP; striped bar) administered 20 minutes before being placed in the runway for DBA/2J mice (n = 13) (b), *Penk*^f/f^ controls (n = 6) (c), or mice lacking enkephalin from D2-MSNs (D2-*Penk*KO, n = 7) (d). Latency to exit the start box was also unchanged on the next day following a saline-prime (right orange bar). **e-h)** Morphine priming acutely increased the time to reach the goal box for *Penk*^f/f^ (g) and D2-*Penk*KO mice (h), but not for DBA/2J mice (f). The latency to reach the goal box decreased following a saline prime administered the day after the morphine prime for DBA/2J, *Penk*^f/f^, and D2-*Penk*KO mice. **i-l.** Morphine priming had no effect on the number of retreats for DBA/2J (j), *Penk*^f/f^ (k), D2-*Penk*KO mice (l). However, the day after the morphine prime, retreats decreased for DBA/2J mice relative to baseline. **m-p)** Morphine priming had no effect on the retreat distance for DBA/2J (n), *Penk*^f/f^ (o), D2-*Penk*KO mice (p), though retreat distance was marginally lower the day after the morphine prime compared to baseline for DBA/2J mice only. **q-t)** The average distance per retreat was not affected by a morphine prime or subsequent saline prime for DBA/2J (r), *Penk*^f/f^ (s), D2-*Penk*KO mice (t). * *p* < 0.05. Data are shown as mean ± SEM. Individual values shown as male (grey) and female (white). Schematics created with Biorender.com.

These data suggest morphine does not alleviate an anxiogenic state and decrease cocaine avoidance. This could indicate that prior reports of decreased retreats for a heroin-cocaine combination may be due to an increased reinforcing value compared to cocaine alone. We therefore speculated that the experience of having morphine on board when cocaine was administered increases the subjective reinforcing value of the goal box. To examine this, we analyzed avoidance behavior the day after the morphine-primed test following a saline prime. DBA/2J mice had fewer retreats (F_1.37, 15.73_ = 4.82, *p* < 0.05; post hoc: baseline vs. saline primed, t_12_ = 2.84, *p* < 0.05) and marginally less retreat distance (F_1.46, 16.76_ = 3.5, *p* = 0.06; post hoc: baseline vs. saline primed, t_12_ = 2.3, *p* = 0.08) on the saline-primed day compared to baseline. Similarly, the latency to goal was lower on the saline-primed day relative to the morphine-primed day (F_1.11, 13.3_ = 6.77, *p* < 0.05; post hoc: baseline vs. saline primed, t_12_ = 3.33, *p* < 0.05). There were no differences in the latency to exit the start box or distance per retreat bout on the saline-primed day compared to baseline. In D2-*Penk*KO and *Penk*^f/f^, the latency to the goal box was lower on the saline-primed day compared to the morphine-primed day (post hoc, t_12_ = 3.27, *p* < 0.05), though no other latency or retreat variables were affected. These data suggest a morphine prime does not alleviate an anxiogenic state to self-administer cocaine in the runway. However, morphine may increase the reinforcing value of cocaine on subsequent trials in DBA/2J mice, but not those on a C57BL/6J background.

## Discussion

The negative affective state induced by falling brain levels of cocaine or acute cocaine administration is a commonly self-reported phenomenon and has been inferred in rats through ultrasonic vocalizations and the development of an approach-avoidance conflict to self-administer cocaine. In the current study, we sought to reproduce cocaine approach-avoidance in a mouse model of the self-administration runway and then used it to investigate how the balance between striatal dynorphin and enkephalin contribute to the development of cocaine avoidance. We observed that mice indeed develop an approach-avoidance conflict similar to reports in rats (Ettenberg and Geist 1993; Geist and Ettenberg 1997; Ettenberg 2004; Jhou et al. 2013; Li et al. 2021; Parrilla-Carrero et al. 2021), and the nature of the expression of this avoidance is strain-dependent. Several reports suggest that the aversive aspect of cocaine withdrawal is associated with dynorphin-mediated KOR signaling in the striatum (Beardsley et al. 2005; Carey et al. 2007; Valdez et al. 2007; Redila and Chavkin 2008), and the aversive state following cocaine administration is associated with lower relative striatal *Pdyn* to *Penk* (Nicot et al. 2025). Despite this, we found no differences in the development of cocaine avoidance in the runway between the DBA/2J and C57BL/6J inbred mouse strains, which are known to have opposing dynorphin-to-enkephalin milieus within the striatum. Moreover, transgenic mice with a pre-existing imbalance between striatal *Pdyn* and *Penk* expression did not show impaired development of cocaine avoidance. Although our results suggest that broad alterations in the striatal balance between dynorphin and enkephalin do not affect cocaine avoidance in the runway, they also emphasize the generalizability of this model across several mouse strains and highlight subtle differences in the expression of cocaine avoidance between the DBA/2J and C57BL/6J strains.

For decades, the DBA/2J and C57BL/6J mouse strains have been contrasted with respect to their addiction-like phenotypes. While initial studies suggested DBA/2J mice are generally less sensitive to the rewarding effects of drugs and alcohol than C57BL/6J mice (Belknap et al. 1993; McGlacken et al. 1995; Risinger et al. 1998; Mittleman et al. 2003), continued work in this field has revealed a more nuanced view. For instance, although DBA/2J mice are teetotalers when it comes to drinking alcohol (Crabbe 2002), they readily self-administer intragastric delivered alcohol, which bypasses the aversive orosensory effects (Fidler et al. 2011). Additionally, compared to C57BL/6J mice, DBA/2J mice acquire cocaine self-administration faster, despite having a lower self-administration rate (Rocha et al. 1998). Considering DBA/2J mice show evidence of higher trait anxiety compared to C57BL/6J (Filiou et al. 2021), it is tempting to speculate that their faster cocaine acquisition is due to a negative reinforcement mechanism, whereby cocaine mitigates their basal anxiety. Furthermore, because DBA/2J mice have greater striatal dynorphin and more KOR binding sites than the C57BL/6J strain, this supposition is consistent with the evidence that dynorphin signaling in cocaine withdrawal drives cocaine seeking (Beardsley et al. 2005; Carey et al. 2007; Valdez et al. 2007; Redila and Chavkin 2008). Accordingly, we expected DBA/2J mice to be more resistant to developing cocaine avoidance and show fewer retreats than C57BL/6J mice. Although cocaine-trained DBA/2J mice had very few retreats, they developed greater latencies to exit the start box and reach the goal box over training, especially in comparison to C57BL/6J mice. This may reflect the inherent higher anxiety levels in DBA/2J mice. In contrast, C57BL/6J exhibited the more “typical” behavior of increasing retreats across trials indicative of cocaine avoidance. Thus, despite the differences in cocaine reinforcement and basal anxiety between DBA/2J and C57BL/6J strains, both express cocaine avoidance, albeit slightly differently from one another and rats. This may suggest striatal opioid peptides do not drive cocaine avoidance but instead underlie the subtle differences in how avoidance is expressed. This interpretation, though, is not consistent with our findings from the *Penk*^f/f^ and D2-*Penk*KO mice. Since this transgenic line is congenic on a C57BL/6J background, we might predict *Penk*^f/f^ to be more “C57BL/6J-like”. Likewise, we might expect D2-*Penk*KO mice to be more “DBA/2J-like” given their higher relative striatal dynorphin-to-enkephalin ratio. Neither strain, however, followed this predicted pattern of avoidance.

Despite the lack of major differences in avoidance between DBA/2J and C57BL/6J mice, this did not completely eliminate a potential role for striatal opioid peptides in cocaine avoidance because these strains have wide genetic differences. Thus, we tested a transgenic mouse line selectively lacking enkephalin from striatal neurons to directly address how striatal enkephalin and the dynorphin-enkephalin balance contributes to cocaine avoidance. Selective deletion of enkephalin from striatal MSNs, however, failed to significantly alter the development of cocaine avoidance. D2-*Penk*KOs and littermate controls developed progressively more retreats over training and increased their latency to reach the goal box, indicating striatal enkephalin is not necessary for the development of cocaine avoidance. Furthermore, it suggests that higher relative dynorphin to enkephalin tone in the striatum does not bias towards the development of cocaine approach over avoidance. This is incongruent with our previous findings showing D2-*Penk*KOs are more resistant to developing cocaine place avoidance in a trace conditioning paradigm (Nicot et al. 2025). One potential explanation for this discrepancy is that cocaine avoidance may arise from distinct sources of cocaine-associated negative affect in the runway compared to the trace conditioning procedure. In trace conditioning, subjects receive a drug injection upon removal from the conditioning chamber. This is thought to imbue the prior environment with the transient negative affective state induced by drug administration before the reinforcing effects begin (Fudala and Iwamoto 1987, 1990; Cunningham and Okorn 1997; Nicot et al. 2025). In contrast, the development of cocaine avoidance in the runway likely results from a combination of cocaine’s transient anxiogenic qualities and the negative affective state brought on by falling brain levels of cocaine. The onset of this negative affective state associated with the cocaine “crash” follows cocaine’s initial reinforcing effect and is consistent with the Opponent-Process Theory of motivation (Solomon and Corbit 1974). Extensive work by Ettenberg and colleagues further supports the premise that the cocaine approach-avoidance conflict exhibited in the runway arises from opposing affective states. For instance, placing rats in an environment 15 minutes after intravenous cocaine administration conditions place avoidance (Ettenberg et al. 1999; Knackstedt et al. 2002), and this timepoint is associated with increased activity of lateral habenula neurons, a region known to encode aversive stimuli (Jhou et al. 2013). Accordingly, pharmacokinetic data indicate cocaine brain levels are likely rapidly falling 15 minutes after intravenous cocaine infusion (Ma et al. 1999). Together, these data suggest subjects experience a transient anxiogenic effect, followed by a rewarding sensation, and ending with an aversive “crash”. Moreover, the negative affective state associated with the cocaine crash is thought to intensify over successive drug use, thereby driving the development of progressive cocaine avoidance in the runway (Ettenberg 2004); however, it is unclear whether the transient anxiogenic effect similarly intensifies over time or if it abates.

Despite evidence that the cocaine crash contributes to development of avoidance in the runway, a cocaine-prime did not alter cocaine avoidance. This may suggest that the more prolonged cocaine withdrawal which occurs over approximately 24 hours is distinct from the more transient “crash” and not a major contributor to cocaine avoidance in the runway. In contrast, pretreatment with the anxiolytic diazepam reduces retreats in the runway (Ettenberg and Geist 1991), and rats allowed to drink ethanol after receiving intravenous cocaine in the goal box also reduce their retreat behaviors (Knackstedt and Ettenberg 2005). Similarly, rats receiving a heroin-cocaine combination in the goal box have fewer retreats compared to rats receiving cocaine alone (Guzman and Ettenberg 2004). Based on this, we predicted pretreatment with morphine would mitigate retreats to the cocaine-paired goal box. Surprisingly, a morphine prime failed to immediately affect cocaine avoidance, indicating morphine does not acutely alleviate the anxiogenic state that develops over training. Although the morphine dose tested (3 mg/kg) showed rewarding efficacy in DBA/2J and C57BL/6J mice (Cunningham et al. 1992), other doses may have distinct effects on cocaine avoidance and should be tested in the future. When tested the next day following a saline-prime, however, retreat behaviors decreased for DBA/2J mice. Thus, we suspect the positive reinforcing qualities of morphine likely combined with those of cocaine. This may also have masked the cocaine-induced negative affective state, which then updated the value of the goal box as more positive and subsequently reduced retreats on the following day.

Taken together, this study indicates that, similar to rats, mice are sensitive to the negative affective states induced by cocaine, and they express this through the development of avoidance behaviors in the self-administration runway. This effect was observed across three different mouse lines, emphasizing the generalizability of these findings; however, future studies should attend to the subtle differences in how avoidance is expressed when testing different mouse lines. Similarly, our findings suggest the ability of morphine to affect cocaine avoidance may differ based on genetic background. Despite finding very few sex differences in DBA/2J mice and none in C57BL/6J mice, conclusions for the D2-*Penk*KO line are limited since we were underpowered to detect sex differences. Additionally, the food-trained control group was not matched for surgical experience or the drug tether. It is possible this contributed to their faster runtimes and fewer retreats, though it is unlikely that this is the main reason for differences between the reinforcer groups.

In conclusion, substantial literature indicates that enkephalins and dynorphins in the striatum play an important role in mediating cocaine seeking behaviors. Our findings suggest that broad changes to the dynorphin/enkephalin milieu within the striatum do not alter the development of cocaine avoidance in the runway. However, it is possible that the role of striatal opioid peptide signaling is more nuanced, and future studies employing fluorescent sensors for dynorphin or enkephalins (Dong et al. 2024) may reveal novel insights into how these peptides modulate cocaine approach and avoidance in real-time.

## Acknowledgements

We would like to thank Jarod Horn for constructing and repairing the self-administration runway and for coding the LabView software. We are grateful to Dr. Andreas Zimmer for providing the *Penk*^f/f^ mouse line.

