## Supplemental Figures for "C57 and DBA mouse strains express distinct cocaine avoidance phenotypes in an operant runway independent of differences in striatal dynorphin-enkephalin balance"

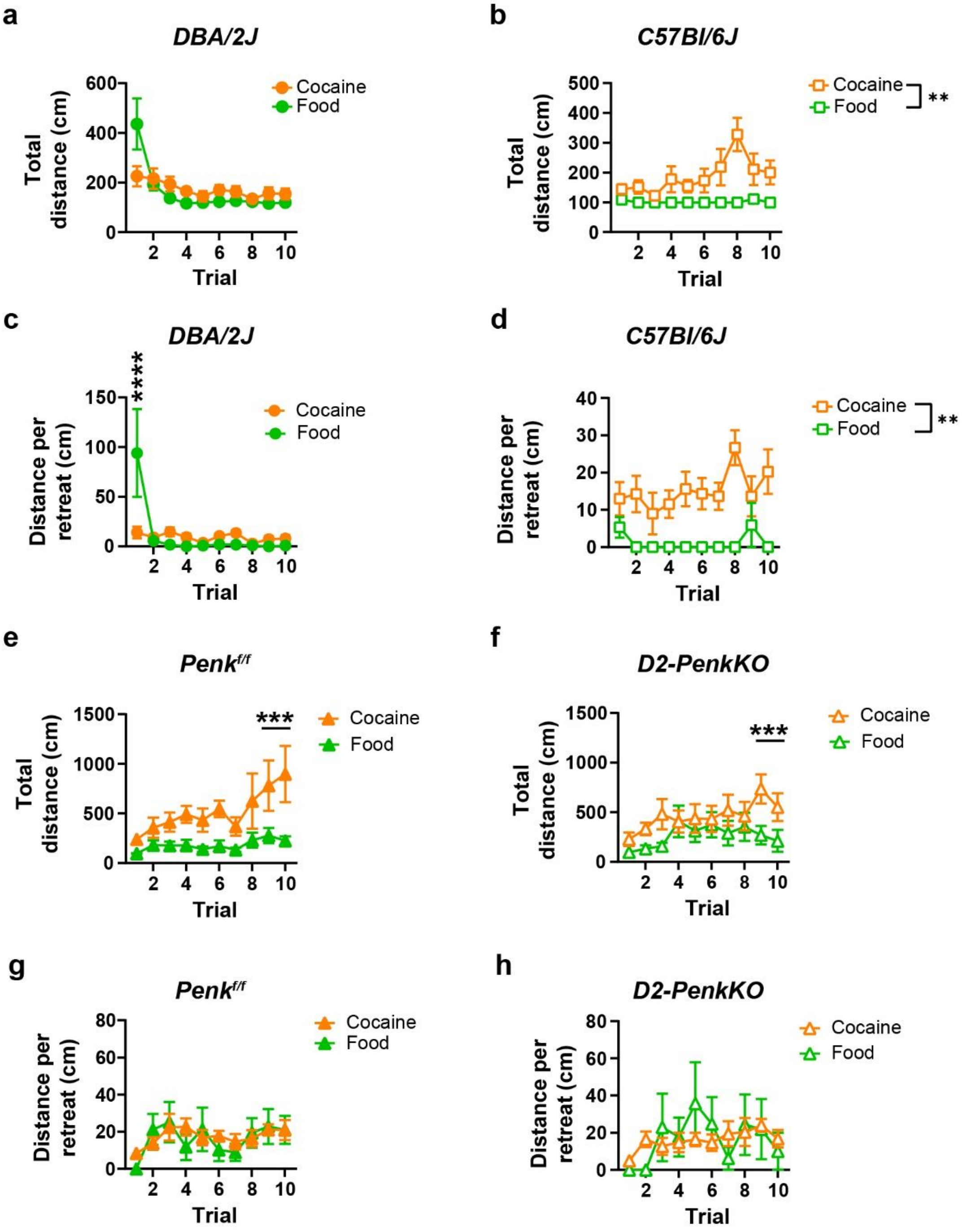


**Fig. S1**

**Locomotor activity and distance traveled per retreat bout across training trials for cocaine and food reinforced DBA/2J, C57BL/6J, *Penk*$^{f/f}$, and D2-*Penk*KO mice. a)** Total distance traveled decreased across training trials but did not differ between cocaine-trained (orange; n = 16) and food-trained (green; n = 16) DBA/2J mice. **b)** Cocaine-trained C57BL/6J mice (n = 14) had greater total distance traveled than food-trained counterparts (n = 10). **c)** Food-trained DBA/2J mice had greater distance traveled per retreat bout on trial 1 than cocaine-trained counterparts. **d)**

Cocaine-trained C57BL/6J mice had greater distance traveled per retreat bout compared to food-trained C57BL/6J mice. **e, f)** Total distance traveled by *Penk*$^{f/f}$ (e) and D2-*Penk*KO (f) mice increased across trials for cocaine-trained compared to food-trained counterparts, specifically at trials 9 and 10 (food: *Penk*$^{f/f}$ n = 9, D2-*Penk*KO n = 6; cocaine: *Penk*$^{f/f}$ n = 10, D2-*Penk*KO n = 10). **g, h)** The distance traveled per retreat bout for *Penk*$^{f/f}$ (g) and D2-*Penk*KO (h) mice did not change across trials and was similar between reinforcers. ** $p < 0.01$, *** $p < 0.001$. Data are shown as mean ± SEM.

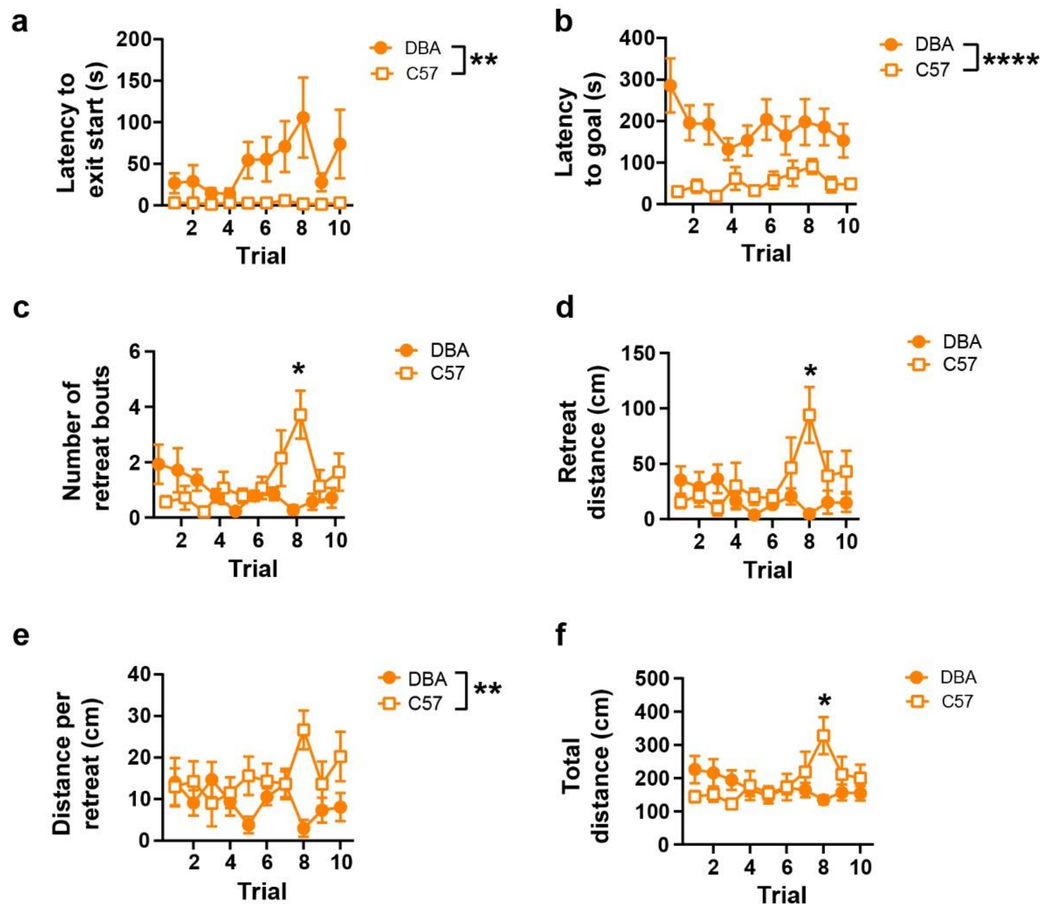


**Fig. S2**

**DBA/2J and C57BL/6J mouse strains express cocaine avoidance differently in the runway.** The measurements of cocaine avoidance during initial training trials for DBA/2J and C57BL/6J are taken from Figures 1 and 2 and replotted to compare the strains with one another. **a, b)** Cocaine-trained DBA/2J mice (filled circles; n = 16) had took longer to exit the start box (a) and to reach the goal box (b) than cocaine-trained C57BL/6J mice (open squares; n = 14). **c, d)** Cocaine-trained C57BL/6J mice developed more retreats (c) and greater retreat distances (d) than DBA/2J mice, specifically at trial 8. **e)** Cocaine-trained C57BL/6J mice had overall greater distance traveled per retreat bout compared to cocaine-trained DBA/2J mice. **f)** Cocaine-trained C57BL/6J mice had nominally greater locomotor activity at trial 8 compared to cocaine-trained DBA/2J mice. # $p = 0.07$, * $p < 0.05$, ** $p < 0.01$, **** $p < 0.0001$. Data are shown as mean ± SEM.

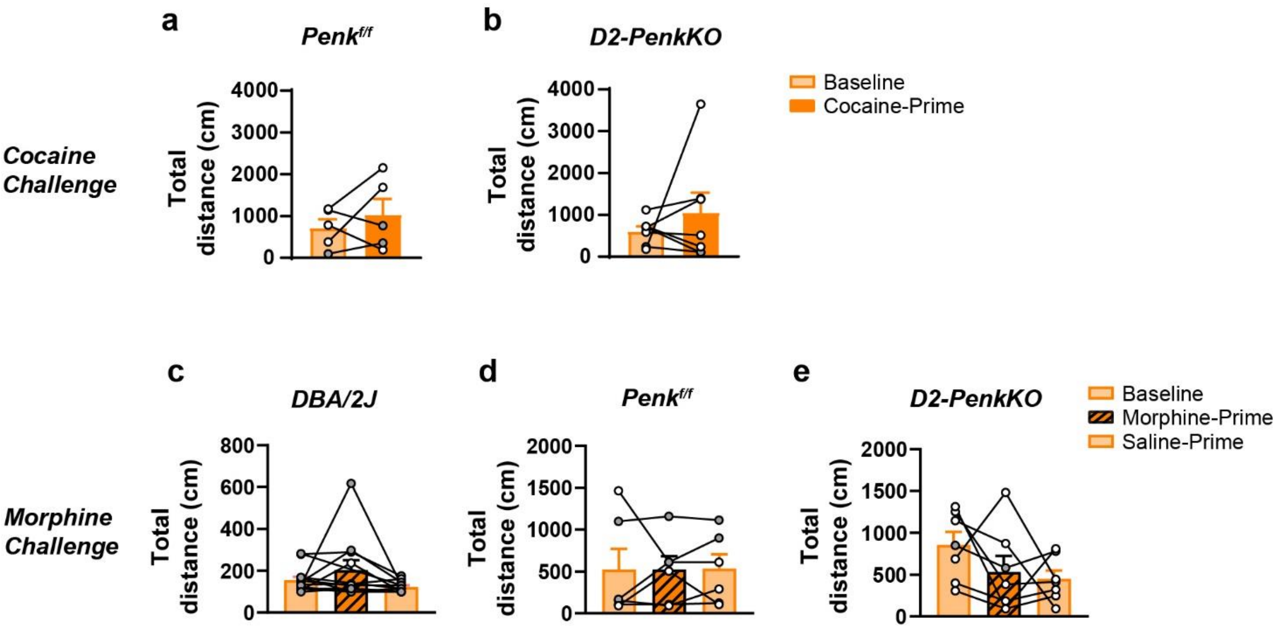


**Fig. S3**

**Locomotor activity following cocaine-prime or morphine-prime does not change. a-c)** Total distance traveled at baseline (average of last 2 training days; light orange) and following a cocaine prime (1.5 mg/kg, IV; dark orange) is shown for DBA/2J (n = 5) (a), *Penk*$^{f/f}$ (n = 5) (b), and D2-*Penk*KO (n = 7) (c) mice. **d-f)** Morphine prime (3 mg/kg, IP, striped bar) 20 minutes before being placed in the runway did not alter locomotor activity for DBA/2J (n = 16) (d), *Penk*$^{f/f}$ (n = 6) (e), or D2-*Penk*KO (n = 7) (f) mice. However, male DBA/2J mice were more active across all days relative to females ($F_{1, 14}$ = 6.36, $p$ < 0.05). Data are shown as mean ± SEM. Individual values shown as male (grey) and female (white).
